# Phosphorylation of Ser111 in Rab8a modulates Rabin8 dependent activation by perturbation of side chain interaction networks

**DOI:** 10.1101/670729

**Authors:** Danial Pourjafar-Dehkordi, Sophie Vieweg, Aymelt Itzen, Martin Zacharias

## Abstract

GTPases are key-players in cellular signaling processes. Phosphorylation of Rab proteins, which belong to the Ras superfamily of small GTPases regulating intracellular transport, has recently been implicated in the pathogenesis of Parkinson Disease (PD). For Rab8a, it was shown that serine 111 phosphorylation (pS111) is dependent on the protein kinase PINK1, and that mimicking the phosphorylation at S111 by a serine/glutamate substitution (S111E) impaired Rab8a activation by its cognate nucleotide exchange factor (GEF) Rabin8. Here, we performed comparative Molecular Dynamics and free energy simulations on Rab8a and Rab8a:Rabin8 complexes to elucidate the molecular details on how pS111 and S111E may influence the interaction with Rabin8. The simulations indicate that S111E and pS111 establish an intramolecular interaction with arginine 79 (R79). In the complex, this interaction persists, and therefore perturbs a favorable intermolecular salt-bridge contact between R79 in Rab8a and the acidic aspartate 187 (D187) in Rabin8. Binding free analysis reveals that S111E and pS111, as well as the mutation R79A, in Rab8a drastically reduce the binding affinity to Rabin8. Combining the R79A mutation with S111E or pS111, respectively, nearly diminishes Rab8a-Rabin8 binding. *In vitro* experiments confirm our computational results showing that the nucleotide exchange rates of the respective Rab8a mutants are decreased by >80% in the presence of Rabin8 compared to wild type. In addition to specific insights into how S111 phosphorylation of Rab8a can influence GEF-mediated activation, the simulations demonstrate how side chain modifications in general can allosterically influence the network of surface side chain interactions between binding partners.

## Introduction

The Rab subfamily of small GTPases is involved in the spatial and temporal regulation of of vesicular trafficking [1–4]. The subfamily consists of ∼60 Rab proteins in humans with specific intracellular localization to mediate signaling functions [5]. The basis for mediating signaling processes is a molecular switch between an inactive guanosine diphosphate (GDP) bound and an active guanosine triphosphate (GTP) bound state. In order to switch from the inactive to the active state, the binding of and activation by guanine exchange factors (GEFs) is required [1]. GEFs are enzymes that stimulate the release of GDP and binding of free GTP to Rab proteins. Additional partner proteins, referred to as effectors, can recognize the active GTP form and promote cellular downstream processes. GTPase activating proteins (GAPs) eventually return Rabs back to their inactive state by stimulating the intrinsic GTPase activity [6, 7]. The Rab activity state is communicated to regulatory proteins and downstream interaction partners by two functionally important loop regions referred to as switch I and II. These regions are conformationally flexible in the inactive state but they become structurally ordered in the active form. Due to their pivotal role in binding to interaction partners, any changes in the conformation of these regions is going to influence the binding profile of the GTPase.

The activity of Rab GTPases can be further modulated by post-translational modifications (PTMs) such as phosphorylation [8–10]. For example, the PTEN-induced kinase 1 (PINK1) is a protein kinase that is indirectly involved in the phosphorylation of a conserved Ser111 residue in the Rab8a, 8b and 13 GTPases *in vivo* [8]. PINK1 is important for mitochondrial quality control, and mutations in PINK1 are associated with autosomal recessive Parkinson’s disease [8]. Recently, it was found that mimicking the phosphorylation of Ser111 by introducing a Ser111->Glu (S111E^Rab8a^) substitution significantly impairs Rab8a activation by its cognate GEF Rabin8 [8].

The crystal structure of GDP-bound Rab8a in complex with Rabin8 provided molecular insights into the mode of this Rab-GEF-interaction [3]. Interestingly, in this 3D structure the S111^Rab8a^ is not directly interacting with Rabin8 (i.e. is not part of the interface). Hence, the decrease on the rate of Rab8a activation by Rabin8 due to S111^Rab8a^-phosphorylation cannot be readily explained by a PTM-induced obstruction of the protein-protein interface. However, S111^Rab8a^ is localized opposite to a negative surface patch of the Rabin8 (see Figure 1). Therefore, the repulsion of charges between the phosphorylation-mimicking S111E^Rab8a^ mutation or the Ser111^Rab8a^-phosphorylation and the negative surface patch of Rabin8 may provide a molecular cause for the less efficient complex formation.

**Figure 1.**
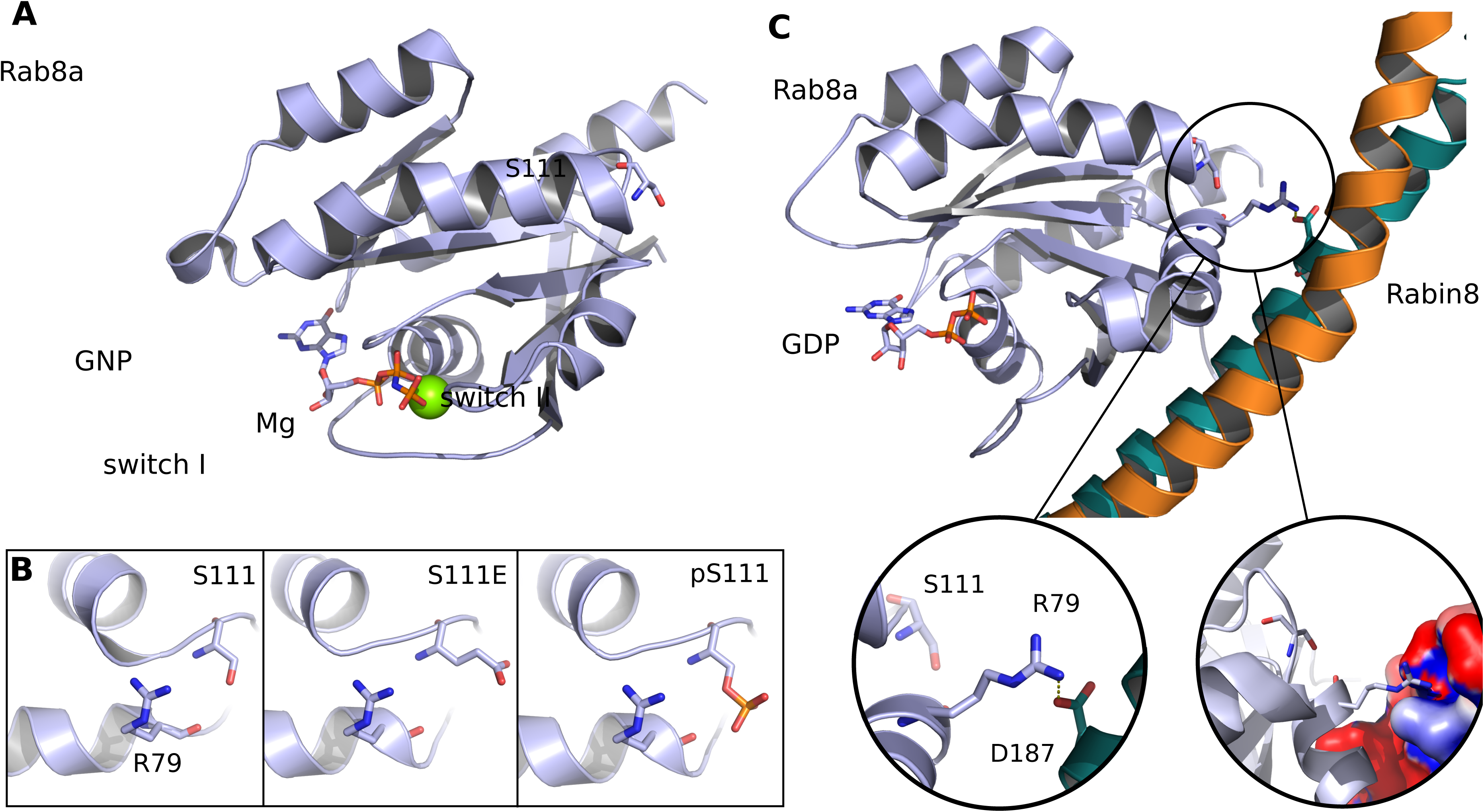
(A) Cartoon representation of the crystal structure of Rab8a bound to a GTP analog (PDB: 4lhw), which served as the starting structure for Rab8a simulations (GTP as atom-color-coded stick model, Mg^2+^ as green van der Waals sphere). (B) Sequence variants (start structures) were generated by replacing S111^Rab8a^ with glutamic acid (S111E^Rab8a^) or phosphoserine (pS111^Rab8a^) *in silico*. (C) Cartoon representation of the crystal structure of the Rab8a:Rabin8 complex and location of bound GDP and of residues S111^Rab8a^, R79^Rab8a^ and D187^Rabin8^ as sticks. The insets illustrate the contact of R79^Rab8a^ and D187^Rabin8^ and the negative patch (red) of Rabin8’s electrostatic surface around residue D187^Rabin8^, respectively.

In the present study we investigate the Rab8a:Rabin8 complex using Molecular Dynamics (MD) simulations to elucidate the molecular mechanism by which the Ser111^Rab8a^-phosphorylation impairs the interaction with Rabin8. We compare the Rabin8-binding of wild type Rab8a with the corresponding S111E^Rab8a^-mutant and pS111^Rab8a^ –modified form. In the complex, D187^Rabin8^ interacts with the switch II-residue R79^Rab8a^ to form a favorable salt-bridge interaction stabilizing the complex. However, in case of the Ser111-phosphorylation or S111E substitution, we identified intramolecular side chain interactions in Rab8a between S111E^Rab8a^/pS111^Rab8a^ and R79^Rab8a^. This interaction weakens or even disrupts the interaction with D187^Rabin8^ in the complex with Rabin8. The simulations demonstrate that R79^Rab8a^ plays a key role in mediating polar interactions between Rab8a and Rabin8 that can be perturbed by introducing the S111E^Rab8a^ mutation or upon S111^Rab8a^ phosphorylation. The simulation results could be confirmed by experimental *in vitro* measurements showing that the Rabin8-mediated nucleotide exchange rate of Rab8a variants (S111E^Rab8a^, R79A^Rab8a^) is decreased by >80% compared to wild type Rab8a.

The study gives insights into the molecular mechanism of signaling modulation by phosphorylation of Rab8a. It furthermore is a model system on how modifications of polar and charged residues adjacent to (but not localized in) a protein-protein interface can allosterically modulate binding strength.

## Results and Discussion

In a previous study it has been found that the phosphorylation mimicking S111E^Rab8a^ substitution (and possibly also S111^Rab8a^ phosphorylation) in Rab8a impairs the activation by its cognate GEF Rabin8 [8]. In order to investigate the molecular origin of this effect we performed a series of Molecular Dynamics (MD) simulations of isolated Rab8a variants and in complex with Rabin8, starting from the known structures (Figure 1). The substitutions were performed *in silico* to yield Rab8a S111E^Rab8a^ and pS111^Rab8a^ variants, both in isolated Rab8a and in the complex with Rabin8 (see Methods). As a first step we performed simulations on the isolated Rab8a variants (in complex with GTP or GDP) starting from the experimental structure in complex with a bound GTP analog (PDB: 4lhw) [3].

On the simulation time scale of 200 ns, the sampled states remained close to the start structure with a similar overall root-mean-square-deviation (RMSD) relative to the start structure of all variants (Figure 2). On this time scale, fluctuations of the switch I and II regions (in particular for the GDP-bound cases) but no unfolding processes were observed (Figure 2, Supporting Information Material Figure S1). However, both the phosphorylation mimetic S111E^Rab8a^ as well as the pS111^Rab8a^ variant formed transient hydrogen bonding states with the side chain of R79^Rab8a^ in the neighboring switch II region (Figure 2). Especially pS111^Rab8a^ formed a robust salt-bridge contact to R79^Raba8^ with two H-bonds for 75 % of the simulation time. In only less than 10% (S111E^Rab8a^) and 2 % (pS111^Raba8^) of the simulation time, there was no contact with R79^Rab8a^. The sampling of these intramolecular H-binding contacts was observed in both the simulations with GTP- and GDP-bound Rab8a (Figure 2). In contrast, no stable contacts between S111^Rab8a^ and R79^Rab8a^ were observed in the MD simulations of wild type Rab8a.

**Figure 2.**
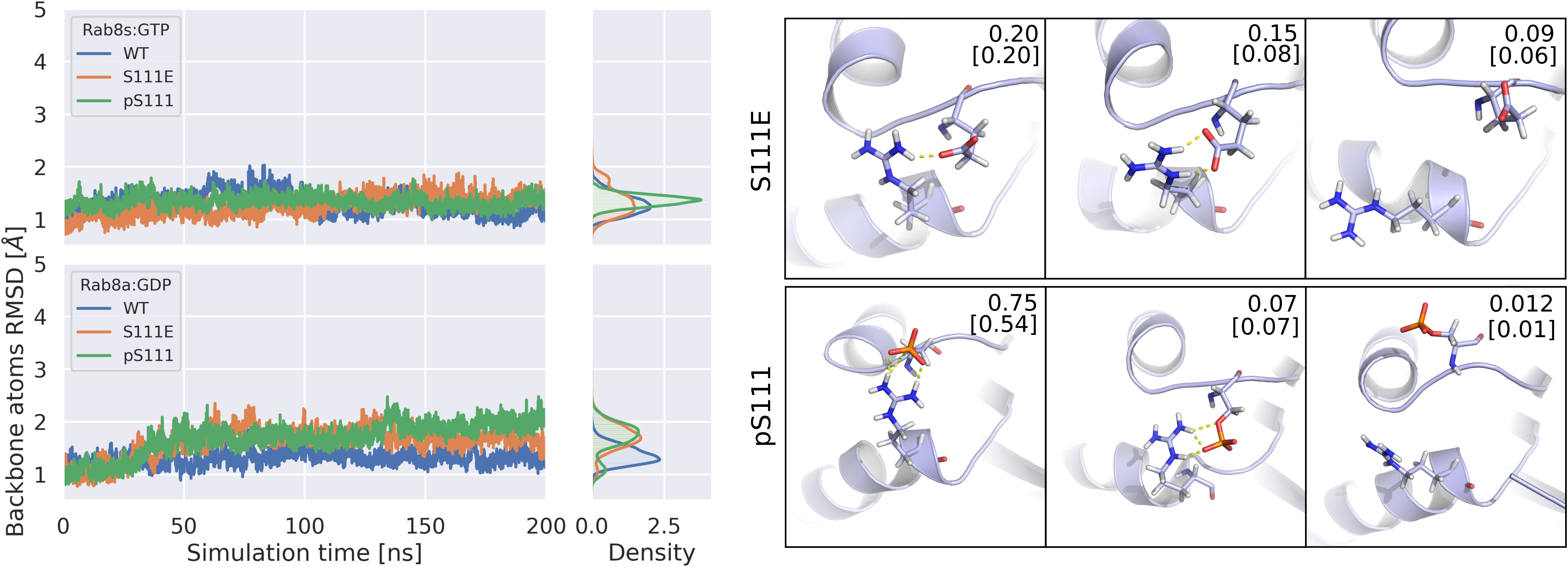
MD-simulations on isolated Rab8a variants in complex with GTP or GDP. Root-mean-square deviation (RMSD) of protein backbone with respect to the initial structure vs. simulation time for wildtype (WT) Rab8a and Rab8a(S111E) and Rab(pS111) variants bound to GTP (upper left panel) and GDP (lower left panel). The RMSD probability density distributions are also indicated. Representative conformations of the most populated 3 clusters (Right panel) obtained from cluster analysis of the trajectories along with their population for GDP-bound Rab8a(S111E) and Rab8a(pS111) cases. The numbers in parentheses indicate the population of the corresponding equivalent clusters in the simulations including bound GTP.

As a next step, MD-simulations of the Rab8a variants in complex with Rabin8 were performed starting from the known crystal structure (GDP-bound, PDB: 4lhy, residue substitution at position S111^Rab8a^ by *in silico* mutation). For all variants, the RMSD with respect to the start structure and the root-mean-square fluctuations (RMSF) (Supporting Information, Figure S2) was similar indicating that the substitutions do not significantly alter the overall structure of the complex.

However, in the case of wild type Rab8a in complex with Rabin8, the R79^Rab8a^ can form a hydrogen bonded salt bridge contact to D187 of Rabin8 (D187^Rabin8^). This hydrogen bonded state was also observed as the dominant local conformational cluster during the simulations of the wild type Rab8a in complex with Rabin8 (Figure 3, 4), suggesting that the contact contributes favorably to the stability of the Rab8a-Rabin8 complex. It is characterized by a short distance between side chain groups of R79^Rab8a^ and D187^Rabin8^ and a larger distance of R79^Rab8a^ and S111^Rab8a^. Indeed, several conformational clusters for the arrangement of the D187^Rabin8^, R79^Rab8a^ and S111^Rab8a^ side chains could be distinguished for the wild type case during the simulations, mostly with direct intermolecular contacts between R79^Rab8a^ and D187^Rabin8^ (Figure 3, Supporting Information Figure S3).

**Figure 3.**
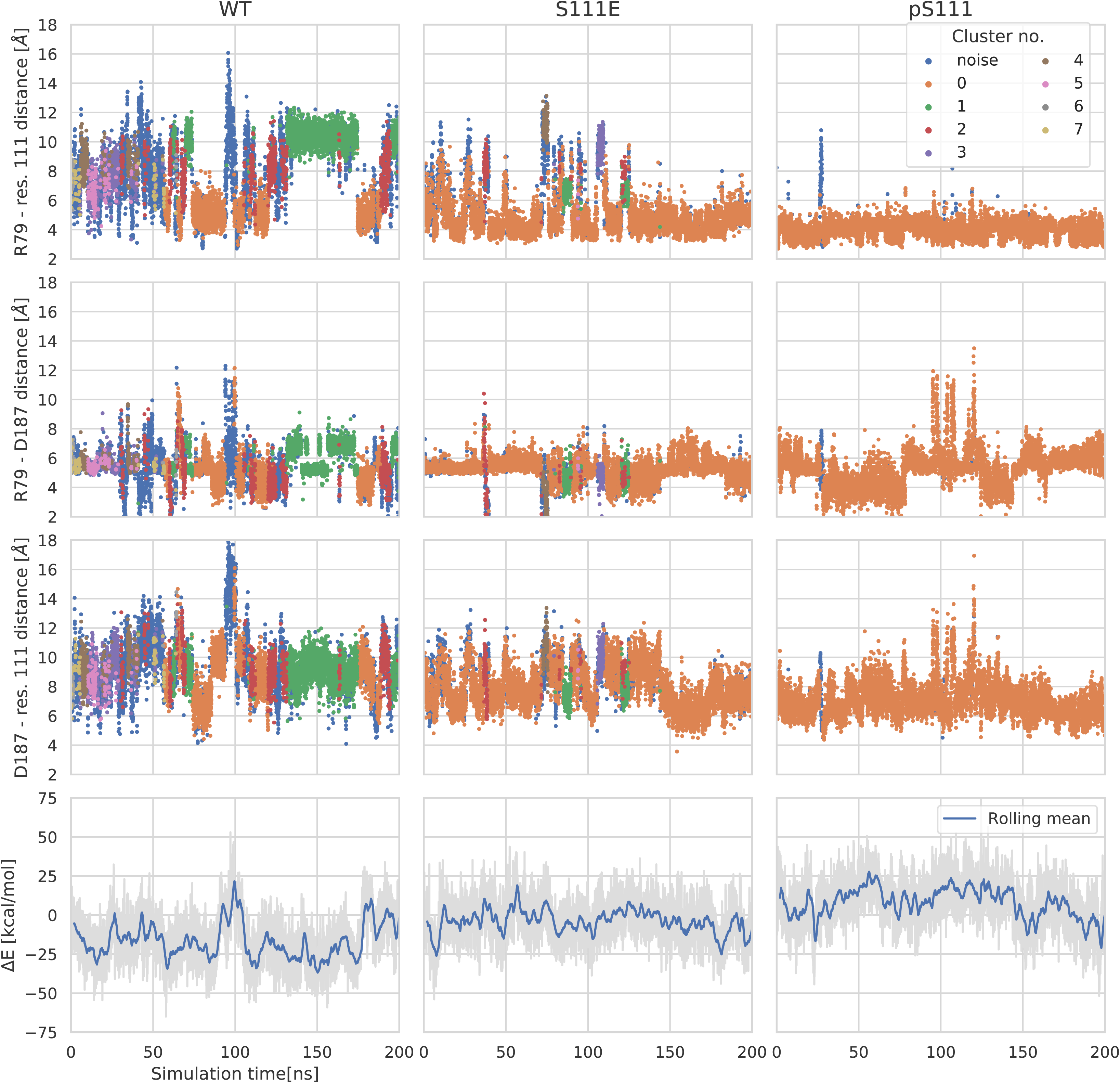
Sampling of conformational substates and interatomic distances involving residues R79^Rab8a^, S111E^Rab8a^, pS111^Rab8a^ and D187^Rabin8^. In the upper panels the assignments to most populated clusters (0-6) are indicated by different point colors. The clusters are numbered based on their population in a descending order (corresponding cluster structures are shown in Supporting Information, Figure S3). The intramolecular distances between R79^Rab8a^/D187^Rabin8^ and residue S111^Rab8a^ are calculated between the arginine amino group (NH_2_) or OG of aspartic acid and the side chain OG of S111^Rab8a^, CG of S111E^Rab8a^ and OG of pS111^Rab8a^, respectively. The intermolecular distances between R79^Rab8a^ and D187^Rabin8^ are the hydrogen-bond distances of the arginine amino group and OG of D187^Rabin8^. Rab8a:Rabin8 interaction energies are calculated using the MMPBSA approach. The blue lines in energy plots indicate the mean over a rolling 2 ns window.

**Figure 4.**
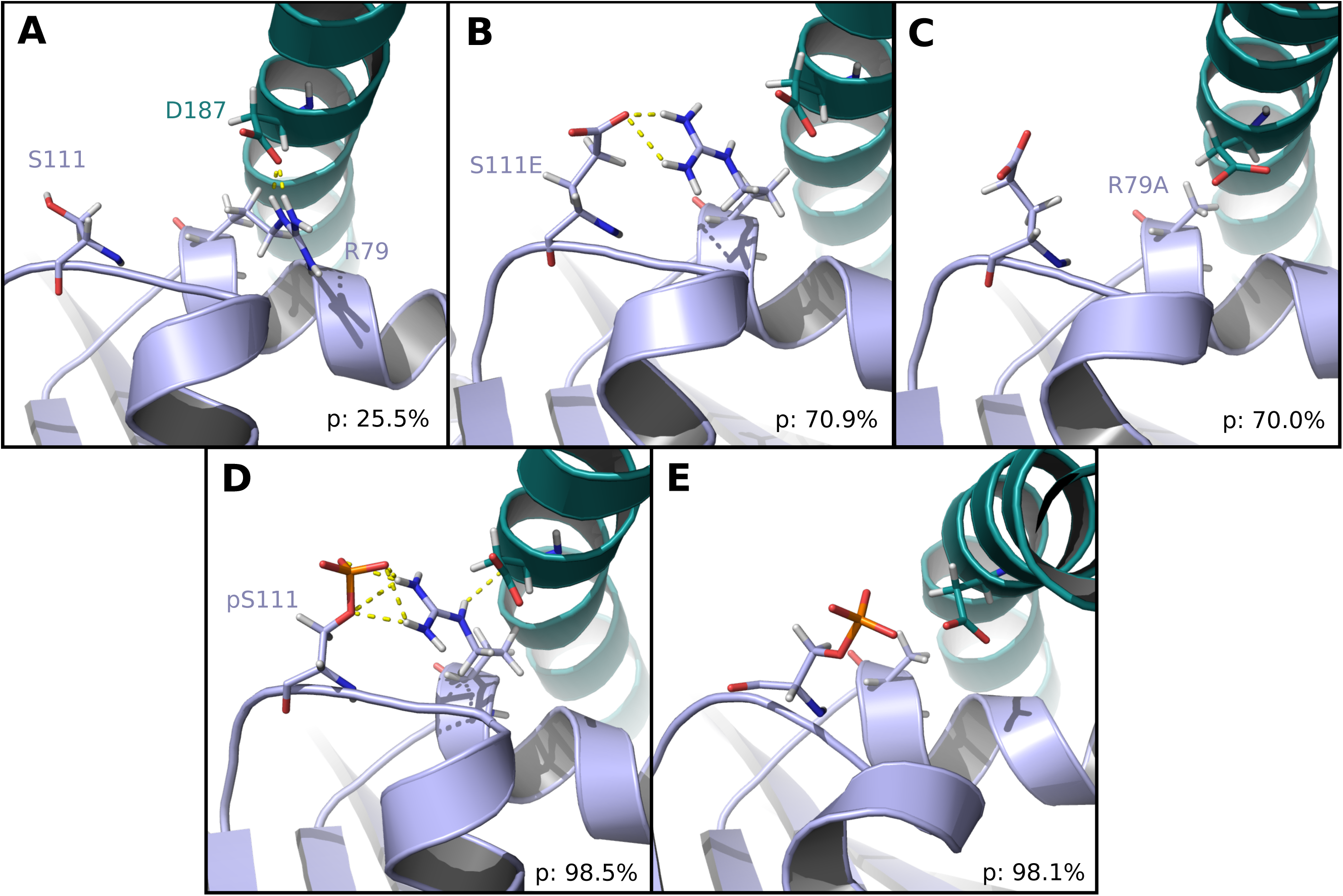
Dominant conformational states observed in simulations of Rab8a:Rabin8 complexes. (A) Representative snapshot of the most populated cluster obtained from 200 ns MD simulations of wild-type Rab8a in complex with Rabin8. (B) Same as (A) but for Rab8a(S111E) in complex with Rabin8. (C) Most populated cluster representative for the MD simulations of the Rab8a(S111E,R79A):Rabin8 complex. (D) representative snapshot of the Rab8a (pS111):Rabin8 complex. (E) Same as in (D) but for the Rab8a (pS111,R79A):Rabin8 complex. Hydrogen bonds are illustrated with yellow dashes. The numbers on the lower right in each panel indicate the population of the clusters as percentage of the frames.

In both cases of pS111^Rab8a^ modification and S111E^Rab8a^ substitution, the simulations indicate that the intramolecular contact to R79^Rab8a^ persists in the simulations in complex with Rabin8 (Figure 3). The average distance of residue S111^Rab8a^ to R79^Rab8a^ is considerably shorter compared to the wild type case (WT-complex: 7.8 Å, Rab8a(S111E)-complex: 5.1 Å, Rab8a(pS111)–complex: 4.1 Å). In the most dominantly sampled states, the R79^Rab8a^-D187^Rabin8^ contact is disrupted (or shows only a weak H-bonding geometry, Figure 4) and some sampled states allow for simultaneous contacts of all three residues (such that the average R79^Rab8a^-D187^Rabin8^ distance is similar in all simulations of the complexes, WT-complex: 5.4 Å, Rab8a(S111E)-complex: 5.2 Å, Rab8a(pS111)–complex: 5.3 Å). The cluster analysis of the arrangement of the three residues in the simulations of the S111E^Rab8a^ and pS111^Rab8a^ variants indicated sampling of fewer distinct clusters compared to the wild type (Figure 3, Supporting Information, Figure S3). Interestingly, due to the formation of the intramolecular contact between residue S111^Rab8a^ and R79^Rab8a^, the average distance of residue S111^Rab8a^ relative to D187^Rabin8^ is also reduced in the simulations of the pS111^Rab8a^ and S111E^Rab8a^ variants (WT-complex: 9.2 Å, Rab8a(S111E)–complex: 8.0 Å, Rab8a(pS111)– complex: 7.1 Å). It brings the negative charges of these residues closer together, and thus is expected to weaken the binding (see below). Finally, control simulations of the double substitution Rab8a(S111E, R79A) or Rab8a(pS111, R79A) in complex with Rabin8 indicate no stable arrangement with close contacts of the side chains R79^Rab8a^, S111^Rab8a^ and D187^Rabin8^ (snapshots of most dominant states are indicated in Figure 4C,E).

In order to quantify the effect of the S111E^Rab8a^ and pS111^Rab8a^ substitution in Rab8a on the binding to Rabin8, we evaluated the trajectories using the molecular mechanics-Poisson-Boltzmann surface area (MMPBSA) method [11, 12]. It should be emphasized that these calculations do not include all contributions to binding (they neglect for instance conformational entropy contributions), but allow a semi-quantitative comparison of relative binding affinity of the variants. The assumption is that the reduction in conformational entropy upon complex formation is similar for all variants. The MMPBSA calculations indicate only a small binding affinity difference in favor of Rab8a:GDP vs. Rab8a:GTP to Rabin8, which is in line with the experimental evidence that there is no strong preference for Rabin8 GEF interactions with the GDP vs. GTP bound forms of Rab8a [3]. Both, the substitution S111E^Rab8a^ as well as the pS111^Rab8a^ modification resulted in a calculated reduction of the binding strength (by about ∼9-10 kcal/mol). Apparently, this gives an estimate of the energetic contribution of the favorable R79^Rab8a^-D187^Rabin8^ contact that is frequently sampled in the case of the Rab8a(WT):Rabin8 complex, but either disrupted or weakened by the presence of the nearby negatively charged residues S111E^Rab8a^ or pS111^Rab8a^ in the Rab8a variants. It is important to emphasize that the calculated magnitude of this binding energy reduction is likely an overestimation because the entropic contributions to restrict the conformational freedom of the side chain motion upon binding is not included. Since our MD-simulations indicate that the R79^Rab8a^ residue may play a key role in mediating the effect of the S111^Rab8a^ modification, which does not directly contact the Rabin8 partner, we used the “alanine scan” option in the MMPBSA approach to investigate the effect of substituting the R79^Rab8a^ by alanine (R79A^Rab8a^). Already in case of the wild type in complex with Rabin8, the R79A^Rab8a^ is predicted to significantly reduce the Rab8a:Rabin8 binding affinity (Table 1). In case of the S111E^Rab8a^ and pS111^Rab8a^ variants of Rab8a, a further reduction in affinity was observed. With the loss of contact between R79^Rab8a^ and D187^Rabin8^ by replacing the arginine with alanine (R79A^Rab8a^), an attractive force between the two proteins is eliminated, and by addition of the phospho-mimetic S111E subsitution (S111E-R79A ^Rab8a^), a repulsive electrostatic force between the two negatively charged residues (S111E^Rab8a^ and D187^Rabin8^) is predicted to weaken the complex affinity even more. This is reflected by the sum of Coulomb and polar-solvation contributions that represent the electrostatic contribution to binding. This contribution is more positive (by 5-9 kcal/mol) for all the R79A^Rab8a^ variants compared to the cases with no R79^Rab8a^ mutation (Table 1).

**Table 1.** Calculated binding free energy (in kcal/mol) of Rabin8 in complex with Rab8a variants

| Mean contributions to binding | Rabin8-Rab8a<sub>GDP</sub> | Rabin8-Rab8a (R79A)<sub>GDP</sub> | Rabin8-Rab8a (S111E)<sub>GDP</sub> | Rabin8-Rab8a (S111E,R79A)<sub>GDP</sub> | Rabin8-Rab8a (pS111)<sub>GDP</sub> | Rabin8-Rab8a (pS111,R79A)<sub>GDP</sub> | Rabin8-Rab8a<sub>GTP</sub> |
| --- | --- | --- | --- | --- | --- | --- | --- |
| $\Delta E_{\text{vanderWaals}}$ | -112.9 | -111.0 | -110.5 | -109.2 | -121.6 | -118.9 | -116.9 |
| $\Delta E_{\text{Coulomb}}$ | -648.7 | -480.1 | -460.8 | -288.9 | -407.2 | -245.7 | -535.9 |
| $\Delta E_{\text{polar\_solvation}}$ | 666.3 | 506.1 | 488.0 | 324.8 | 441.6 | 285.1 | 560.6 |
| $\Delta E_{\text{cavity}}$ | -87.5 | -83.8 | -85.9 | -82.7 | -95.6 | -92.7 | -89.7 |
| $\Delta E_{\text{dispersion\_solvent}}$ | 171.3 | 165.9 | 169.2 | 164.6 | 181.2 | 177.0 | 173.3 |
| $\Delta G_{\text{binding}}$ | -11.6±0.3 | -2.9±0.3 | -0.01±0.3 | 8.5±0.3 | -1.8±0.4 | 4.8±0.4 | -8.7±0.4 |
| $\Delta\Delta G_{\text{binding}}$ | - | 8.6±0.4 | 11.5±0.5 | 20.0±0.5 | 9.8±0.5 | 16.4±0.47 | 2.8±0.5 |

Mean binding energy contributions were obtained as MMPBSA averages from MD-simulations (1000 snapshots) of the Rab8a-Rabin8 complexes (see Methods). $\Delta G_{\text{binding}}$ corresponds to the estimated absolute mean binding free energy and $\Delta\Delta G_{\text{binding}}$ represents the relative binding free energy with respect to the wild type Rabin8-Rab8a<sub>GDP</sub> complex. Mean energies and free energies are stated in kcal/mol.

In addition to the average effect of the substitutions we also split the trajectories into sets of frames that belong to different conformational clusters formed by the residues R79^Rab8a^, S111^Rab8a^ and D187^Rabin8^ (see color coding in Figure 3). The MMPBSA analysis of these sets of frames indicates that each cluster of side chain arrangements contributes differently to the binding affinity. Some clusters make a favorable and others a less favorable or even unfavorable contribution to binding. For example, in case of the wild type simulations at around 100 ns mostly conformations with a R79^Rab8a^-D187^Rabin8^ distance > 4 Å are sampled resulting in an overall slightly positive interaction energy (ΔE >0) compared to states with a smaller R79^Rab8a^-D187^Rabin8^ distance (Figure 3). The S111E^Rab8a^ substitution or phosphorylation results in a shift of the clusters to more states with an unfavorable effect on binding. The interaction energy (indicated as ΔE) is more positive compared to the wild type simulations especially in time intervals that correspond to conformations with increased R79^Rab8a^-D187^Rabin8^ distances (illustrated in the last row of panels in Figure 3). The arrangement forms a model system how modifications of residues not being part of an interface can mediate or control binding affinity by changing the network of contacts of the nearby residues (in the present case, of R79^Rab8a^) with residues on the partner protein.

The simulations suggest differences in the binding affinity of the Rab8a variants to Rabin8, which should result in decreased nucleotide exchange rates for those variants compared to wild type Rab8a. With the exception of the pS111^Rab8a^ variant, it was possible to generate, express and purify all Rab8a variants (Supporting Information Figure S4). We determined the kinetics of Rabin8-stimulated nucleotide exchange reactions for Rab8a(WT), Rab8a(S111E), Rab8a(R79A) and Rab8a(S111E-R79A). The Rabin8-simulated GDP→GTP exchange rates were calculated for each Rab8a variant based on the time-dependent change of intrinsic tryptophan fluorescence (Figure 5). We found that the phospho-mimetic S111E^Rab8a^ and the R79A^Rab8a^ mutants resulted in >80% reduction of the nucleotide exchange rate compared to wild type Rab8a. In the case of the S111E/R79A^Rab8a^ double mutant, the nucleotide exchange activity was almost completely abolished (∼95% decrease). Since the mutations are not on the Rabin8 side, the reduced activity must be due to the reduced binding of Rabin8 to Rab8a variants. Hence, the experimental results are in line with the computational prediction that the presence of S111E^Rab8a^ or pS111^Rab8a^ prevents R79^Rab8a^ to engage in interactions with Rabin8, and therefore reduces the binding affinity [13].

**Figure 5.**
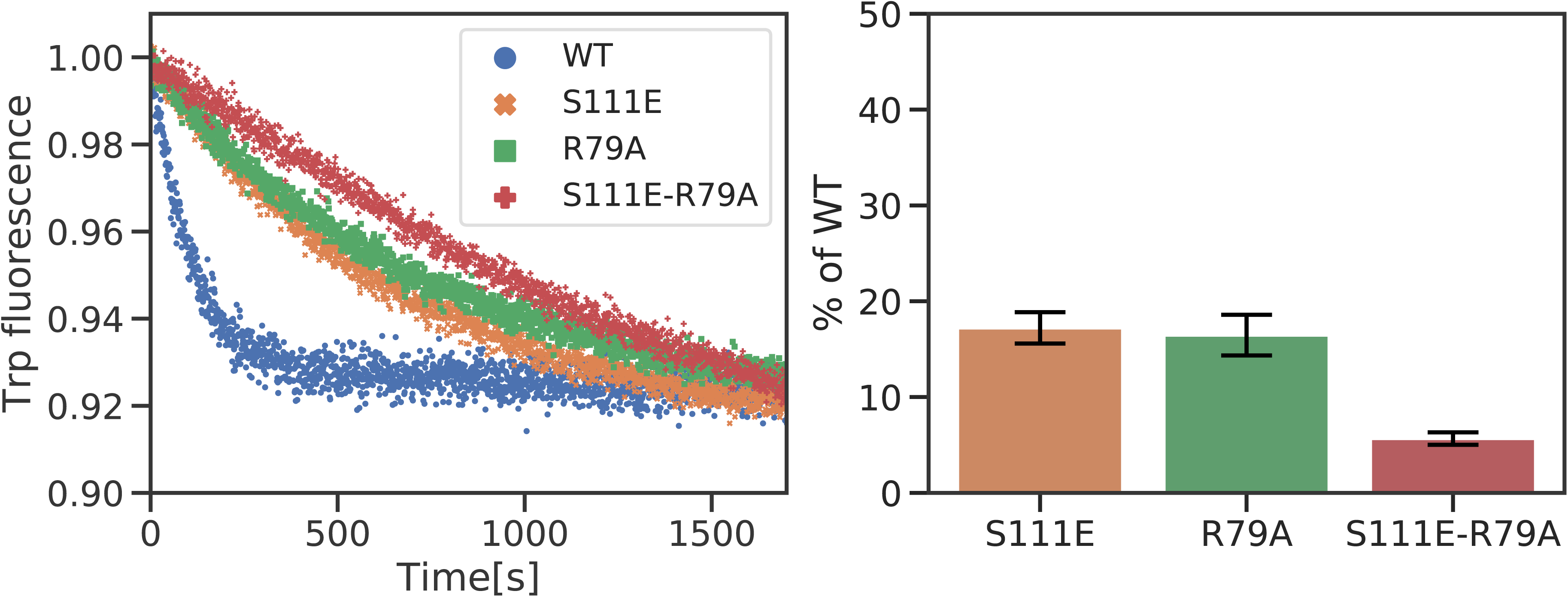
(A) Rabin8-catalyzed GDP-GTP exchange reaction observed as a decrease in tryptophane fluorescence in Rab8a(WT), Rab8a(S111E), Rab8a(R79A) and Rab8a(S111E-R79A). (B) Measured k_obs_ for the exchange reaction of the Rab8a variants in comparison to Rab8a(WT).

## Conclusions

Post-translational modifications may play a role in modulating signaling properties of GTPases [8, 14]. Indeed, for Rab8a it has been previously shown that mimicking the phosphorylation at serine 111 (S111E^Rab8a^) reduces the GEF-mediated activation by Rabin8 [8]. Interestingly, the structure determination of the Rab8a:Rabin8 complex [3, 8] revealed that S111^Rab8a^ is not directly located at the interface to Rabin8, and that even upon phosphorylation no direct contact to Rabin8 is sterically possible. In the present study, we investigated the molecular details on how S111E^Rab8a^ and pS111^Rab8a^ affect the interaction with Rabin8 using a series of MD-simulations and free energy calculations. Both, the *in silico* S111E^Rab8a^ substitution as well as the pS111^Rab8a^ modification tend to form intramolecular salt bridge-like contacts to the nearby R79^Rab8a^ residue within the switch II region. This was observed in the isolated Rab8a variant and in the complex with Rabin8 as dominant conformational states that in case of the complex perturbs and even disrupts the R79^Rab8a^-D187^Rabin8^ salt-bridge contact in contrast to the wild-type complex. The importance of R79^Rab8a^ for stabilizing the binding of Rab8a to Rabin8 could be demonstrated by analyzing the R79A^Rab8a^ substitution as well as double mutations (R79A^Rab8a^-S111E^Rab8a^, R79A^Rab8a^-pS111^Rab8a^). Although the predicted effects on the change in binding free energy due to the substitutions are likely to be larger than the experimental binding free energy changes, the order of the effect of each variant correlates very well with experimental data on the binding to Rabin8 and the GEF efficiency of Rabin8. *In vitro* experiments investigating the Rabin8-mediated nucleotide exchange reactions confirmed the computational results. We show that the exchange rates of all Rab8a (S111E, R79A) variants are decreased by >80% compared to wild type Rab8a, suggesting that the residue R79^Rab8a^ in Rab8a plays an essential role in mediating the binding of Rabin8, and that S111E^Rab8a^ interferes with this function.

It is well known that phosphorylation of protein residues can alter conformational equilibria, and in turn influence binding to protein partners. Examples are the phosphorylation of Arg/Ser rich proteins (RS-proteins) resulting in conformational changes and disruption of binding to RNA molecules [15], or the phosphorylation of Tyr residues in Ras-GTPases that alter the switch I and II conformation directly affecting the interaction with effectors [16, 17]. The present study demonstrates a potential mechanism on how a chemical modification of a residue, which is not part of the binding interface, can modulate signaling events due to altering the neighboring side chain interaction network that is part of the interface with the partner protein. Such mechanism can be of particular importance when it comes to the fine-tuning of cell signaling events, where a complete disruption of the binding, and thus signaling, would be detrimental. The perturbation or alteration of interacting side chain networks can potentially be the basis of allosteric effects mediated not by chemical modification of side chains, but by binding of an allosteric effector adjacent to the interface with another binding partner. Indeed, for HLA-DR (MHC class II) molecules the conformational change of a Trp side chain induced by binding of the co-chaperone HLA-DM (at a site not overlapping with the peptide binding groove) has been found to control peptide binding and exchange [18]. Similar to the present case the allosteric effect is then mediated by perturbation of a side chain interaction network that mediates the interaction with the binding partner.

## Materials and Methods

### Molecular Dynamics Simulations

All isolated Rab8a simulations started from the crystal structure bound to phosphoaminophosphonic acid guanylate ester (GNP) (PDB: 4lhw) [3]. The Rabin8-bound simulations started from the crystal structure of Rab8a:Rabin8 complex (PDB: 4lhy) [3]. The nitrogen atom between the β- and γ-phosphate in GNP was exchanged with an oxygen atom to model the Rab8a:GTP structure. The terminal phosphate was removed to form an initial model of the Rab8a:GDP complex. In the model of the phosphorylated complex, the Ser111 (S111^Rab8a^) was replaced with phosphoserine (Ser111:pS111^Rab8a^) to form the phosphorylated Rab8 structures. The Amber ff14sb force field [19] was used for the proteins, additional force field parameters for GDP, GTP and phosphoserine were taken from the Amber parameter database at their fully unprotonated states [20, 21]. A recent study by Mann et al. [22] supports the unprotonated state of GTP as the most populated state at neutral pH. For Rab8a:GDP simulations, a Mg^2+^ ion with two bound water molecules was placed next to GDP-phosphate. Using sodium and chloride ions, the salt concentration was adjusted to 0.1 M and the systems were solvated with the TIP3P water model [23, 24]. The solvated systems were equilibrated by a first energy minimization (5000 steps), followed by 25 ps of heating and 50 ps of density equilibration, followed by a simulation in NPT ensemble at 300 K. During these equilibration phases, all protein nucleotide heavy atoms as well as magnesium ions were restraint with a harmonic potential at force constant of 5.0 kcal mol^-1^Å^-2^. Data gathering production simulations were performed without any restraints. The pmemd version of the Amber 16 software package [13] in combination with hydrogen mass repartitioning [25] was used which allows a simulation time step of 4 fs. Long range interactions were included using the particle mesh Ewald (PME) method combined with periodic boundary conditions and a 9 Å cut-off for real space non-bonded interactions. Trajectories were processed and analyzed using CPPTRAJ program [13]. The DBSCAN algorithm was used for clustering of the trajectories with a distance cutoff of 1.0 Å of heavy atoms root mean square deviation (RMSD) and a frame interval of 200ps. Figures were generated using PyMol software package [26].

#### Binding affinity calculations

The interaction energies between Rab8a and Rabin8 were calculated using MMPBSA tool [11] of the AMBER software suite. Five production simulations of 2ns length started from the complex of GDP-bound Rab8a and Rabin8 (PDB: 4lhy) at 300K, generating snapshots every 10ps. A similar set of simulations were carried out starting from the same structure but with S111E^Rab8a^ mutated. Using the ‘alanine scan’ feature of MMPBSA, the contribution of R79^Rab8a^ in each case was evaluated. Calculations were carried out on a sum of 1000 frames for each complex at an ion concentration of 0.1M.

#### Expression and purification of Rab8a(6-176aa)

Rab8a-His_6_ proteins (pET19) were co-expressed with GroEL/S (pGro7, Takara) in BL21(DE3) *Escherichia coli* in LB medium at 25°C overnight. The LB medium contained 1mg/mL arabinose for autoinduction of the expression of GroEL/S, while the expression of Rab8a-His6 was induced by the addition of 0.5 mM IPTG at OD_600nm_ = 0.8. Cells were harvested and resuspended in buffer A (50 mM HEPES, 500 mM LiCl, 10 mM imidazole, 1 mM MgCl_2_, 10 µM GDP, 2 mM β-mercaptoethanol, pH 8.0) containing 1 mM PMSF and DNaseI. Cells were lysed on ice by sonication (60 % amplitude, 5 min, pulse 5 s on and 15 sec off), and insoluble cell debris was removed by centrifugation (48254.4 g, 45 min, 4 °C). Rab8-His_6_ proteins were purified from the supernatant by nickel affinity chromatography (5 ml HiTrap™ Chelating HP column, GE Healthcare). Unspecifically bound proteins were removed by a step wash with 5% buffer B (50 mM HEPES, 500 mM LiCl, 500 mM imidazole, 1 mM MgCl_2_, 10 µM GDP, 2 mM β-mercaptoethanol, pH 8.0) and the Rab8a-His_6_ proteins eluted with a linear gradient from 5-60 % buffer B in 100 ml. The collected fractions were analyzed by SDS-PAGE. Fractions containing Rab8-His_6_ were concentrated to 2 ml using Amicon^®^ Ultra Centrifugal Filters (10,000 MWCO, Merck Millipore), and injected into a Superdex 75 16/60 (GE Healthcare) equilibrated with buffer C (20 mM HEPES, 50 mM NaCl, 1 mM MgCl_2_, 10 µM GDP, 1 mM DTT, pH 7.5). Collected fractions were analyzed by sodium dodecylsulfate (SDS) polyacrylamide gel electrophoresis (PAGE). Pure fractions were pooled, concentrated using Amicon^®^ Ultra Centrifugal Filters (10,000 MWCO, Merck Millipore), and frozen in liquid nitrogen. The protein identity was confirmed by liquid chromatography mass spectrometry.

#### Liquid chromatography mass spectrometry (LC-MS)

Protein samples (0.1 mg/ml, 1 µl) were injected with an UltiMate3000^®^ HPLC system (UHPLC^+^ focused, Dionex) into a ProSwift™ RP-4H column (1x 50 mm, Thermo) at a flow rate of 0.7 ml/min. The proteins eluted with a linear gradient of 5-100 % acetonitrile (0.1 % formic acid) in 6 min. The desalted samples were ionized and analyzed by a LCQ fleet™ system (Thermo) combining electro spray ionization (ESI) with an ion trap mass analyzer.

#### GDP-GTP exchange kinetics

The Rab8a proteins, purified in the presence of 10 µM GDP, were diluted in buffer (20 mM HEPES, 50 mM NaCl, 1 mM MgCl_2_, pH 7.5) in a Quartz SUPRASIL cuvette (Hellma Analytics, Germany) to a concentration of 4 µM. The intrinsic tryptophane (Trp) fluorescence was recorded at λ_em_= 348 nm (λ_exc_= 297nm) at 25 °C with a Fluoromax-4 fluorescence spectrometer (HORIBA Jobin Yvon). When the Trp fluorescence signal stabilized, 50 µM GTP and 2 µM Rabin8 were added. Based on the decrease in Trp fluorescence over time, the GDP-GTP exchange rates were calculated for each Rab8a variant using the following exponential equation:

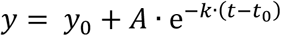

y_0_: final fluorescence value (end point)

A: amplitude of fluorescence change

t: time in seconds

t_0_: initial time point of the reaction

## Acknowledgments

We thank the laboratory of Jason W. Chin (Medical Research Council Laboratory of Molecular Biology, Francis Crick Avenue, Cambridge, CB2 0QH) for providing us the plasmids for genetically encoding phospho-serine and the _serB-BL21(DE3) *E. coli* cells for expression. This work was supported by SFB1035 project grants B02 and B05 funded by the DFG (Deutsche Forschungsgemeinschaft) and by the Michael-J-Fox-Foundation. Computation resources were provided by the Leibniz Supercomputing Centre (LRZ) within grant pr74bi and are highly appreciated. We are grateful to Miratul Muqit for critically commenting on the manuscript.

## Author contributions

MZ conceived the theoretical and computational part of the study, AI conceived the experimental part. All experiments were performed by SV. DPD performed all simulations. All authors analyzed data and wrote the paper.

